# Structural basis of RICs iron donation for iron-sulfur cluster biogenesis

**DOI:** 10.1101/2021.02.18.431928

**Authors:** Liliana S. O. Silva, Pedro M. Matias, Célia V. Romão, Lígia M. Saraiva

**Author notes:** **Corresponding authors:** Lígia M. Saraiva, Instituto de Tecnologia Química e Biológica António Xavier, Universidade Nova de Lisboa, Av. da República, 2780-157 Oeiras, Portugal., Célia V. Romão, Instituto de Tecnologia Química e Biológica António Xavier, Universidade Nova de Lisboa, Av. da República, 2780-157 Oeiras, Portugal.

## Abstract

*Escherichia coli* YtfE is a di-iron protein, of the widespread RIC family, with capacity to donate iron, which is a crucial component of the biogenesis of the ubiquitous family of iron-sulfur proteins. Herein we identify in *E. coli* a previously unrecognized link between the YtfE protein and the major bacterial system for iron-sulfur cluster (ISC) assembly. We show that YtfE establishes protein-protein interactions with the scaffold IscU, where the transient cluster is formed, and the cysteine desulfurase IscS. Moreover, we found that promotion by YtfE of the formation of an Fe-S cluster in IscU requires two glutamates, E125 and E159 in YtfE. Both glutamates form part of the entrance of a protein channel in YtfE that links the di-iron centre to the surface. In particular, E125 is crucial for the exit of iron, as a single mutation to leucine closes the channel rendering YtfE inactive for the build-up of Fe-S clusters. Hence, we provide evidence for the key role of RICs as bacterial iron donor proteins involved in the biogenesis of Fe-S clusters.

**Importance:** The ubiquitous iron-sulfur proteins require specialized cellular machineries to promote the assembly of the cofactor. These systems include proteins that provide sulfur and iron, and scaffold proteins where the cluster is formed. Although largely studied the nature of the iron donor remains to be fully clarified. In this work, we show that *Escherichia coli* YtfE, which belongs to the RIC protein family, establishes protein-protein interactions with two of the major proteins of the ISC system, and we reveal the structural characteristics necessary for the exit of iron ions from YtfE. Altogether our results prove that RICs can be considered a family of iron donor proteins involved in the biogenesis of iron-sulfur containing proteins.

## Introduction

The Repair of Iron Centres proteins (RIC) is a family of di-iron proteins that provide protection to enzymes prone to inactivation by oxidative and nitrosative stresses imposed by the host’s innate immune system (1–3). Firstly discovered in *Escherichia coli*, YtfE is a member of the RIC family present in bacteria, fungi and eukaryotes(1, 4). In particular, homologues of these proteins are encoded in the genomes of a significant number of human pathogens, such as *Bacillus anthracis, Haemophilus influenzae*, and species of the genus *Salmonella, Shewanella, Yersinia* or *Clostridium*. Two RIC paralogues (RIC1 and RIC2) are also present in the eukaryote *Trichomonas vaginalis*, an important human pathogenic protozoan(5). *In vivo* studies showed that RIC deletion in *E. coli* and *S. aureus* generates strains defective in the activity of several iron-sulfur-containing proteins, and RICs contribute to the survival and virulence of *Staphylococcus aureus, Yersinia pseudotuberculosis*, and *Haemophilus influenzae*(1, 6–8).

Bacterial RICs are composed by an N-terminal ScdA-like domain and a C-terminal domain that folds as a four-helix bundle where a histidine/carboxylate di-iron type centre is inserted. In *E. coli* YtfE, the best studied RIC protein, the centre is coordinated by H84, H129, H160, H204, and two μ-carboxylate bridges formed by E133 and E208(9). These residues are highly conserved in the RIC family and are required for the assembly of a stable and functional di-iron cluster(9). Mössbauer and EPR studies showed that the centre of the as-isolated YtfE has properties characteristic of a mixed valence antiferromagnetically coupled Fe(III)–Fe(II) centre with a S=½ ground state(9–11). Mössbauer studies further revealed that one of the iron atoms in the centre is labile, particularly in the mixed-valence Fe(III)–Fe(II) state when compared with the μ-oxo-diferric form Fe(III)-Fe(III)(11). In the reduced form, YtfE Fe(II)–Fe(II) binds NO forming N_2_O(12), which is in line with the intrinsic capacity of di-iron proteins for NO reduction and O_2_ activation/reduction(13). Moreover, in YtfE one of the iron ions of the centre is loosely bound with iron dissociation constants in a range of values that enables the YtfE to donate iron to other proteins(11).

In bacteria, the biogenesis of Fe-S clusters requires specialised machineries, ISC (iron sulfur cluster) being one of the main operative systems. As in many other bacteria, in *E*. coli the first steps of the biogenesis involve IscS, a cysteine desulfurase that catalyses the sulfur mobilisation from cysteine, an iron donor protein, and the scaffold IscU protein where the Fe-S clusters are assembled prior to their transport and insertion into the target apo-proteins(14–19). We previously provided evidences that *E. coli* YtfE acts as an iron donor for the formation of iron-sulfur clusters in IscU(11). In this work, we show that YtfE interacts directly with IscS and IscU. The analysis of the biochemical properties and crystal structures of *E. coli* YtfE wild-type and several site-directed mutants allowed us to identify two glutamates, E125 and E159, with a key role in supporting the ability of YtfE to act as an iron donor protein.

## Results

### YtfE interacts with IscS and IscU for Fe-S cluster biogenesis

We previously reported that *E. coli* YtfE donates iron to form an Fe-S cluster in *E. coli* IscU in a reaction that also required the presence of IscS(11). This observation led us to examine herein whether YtfE interacts with IscS and IscU using a genetic approach, a complementation fluorescence assay and pulldown experiments.

For the genetic approach, we measured the β-galactosidase activities of cells transformed with a pair of plasmids derived from the high-copy number vector pUT18 and the low-copy number vector pKT25(20). These vectors express YtfE, IscS and IscU proteins as fusions to the N- and C-terminal of T25 domain (T25 and T25C, respectively) and to the N- and C-terminal of T18 domain (T18 and T18C, respectively) of the adenylate cyclase enzyme (see Methods). The YtfE-YtfE interaction was used as positive control, as YtfE self-associates forming dimers(2).

We observed the formation of a complex between YtfE and IscS and IscU (Fig. 1a). The interaction between YtfE and IscS was 2-4 times higher in comparison with the control interactions and was dependent on the IscS protein configuration. Concerning the YtfE-IscU interaction, only the conformation in which IscU was expressed from the N-terminal part of the pUT18 fusion protein yielded significant β-galactosidase activity.

**Fig. 1.**
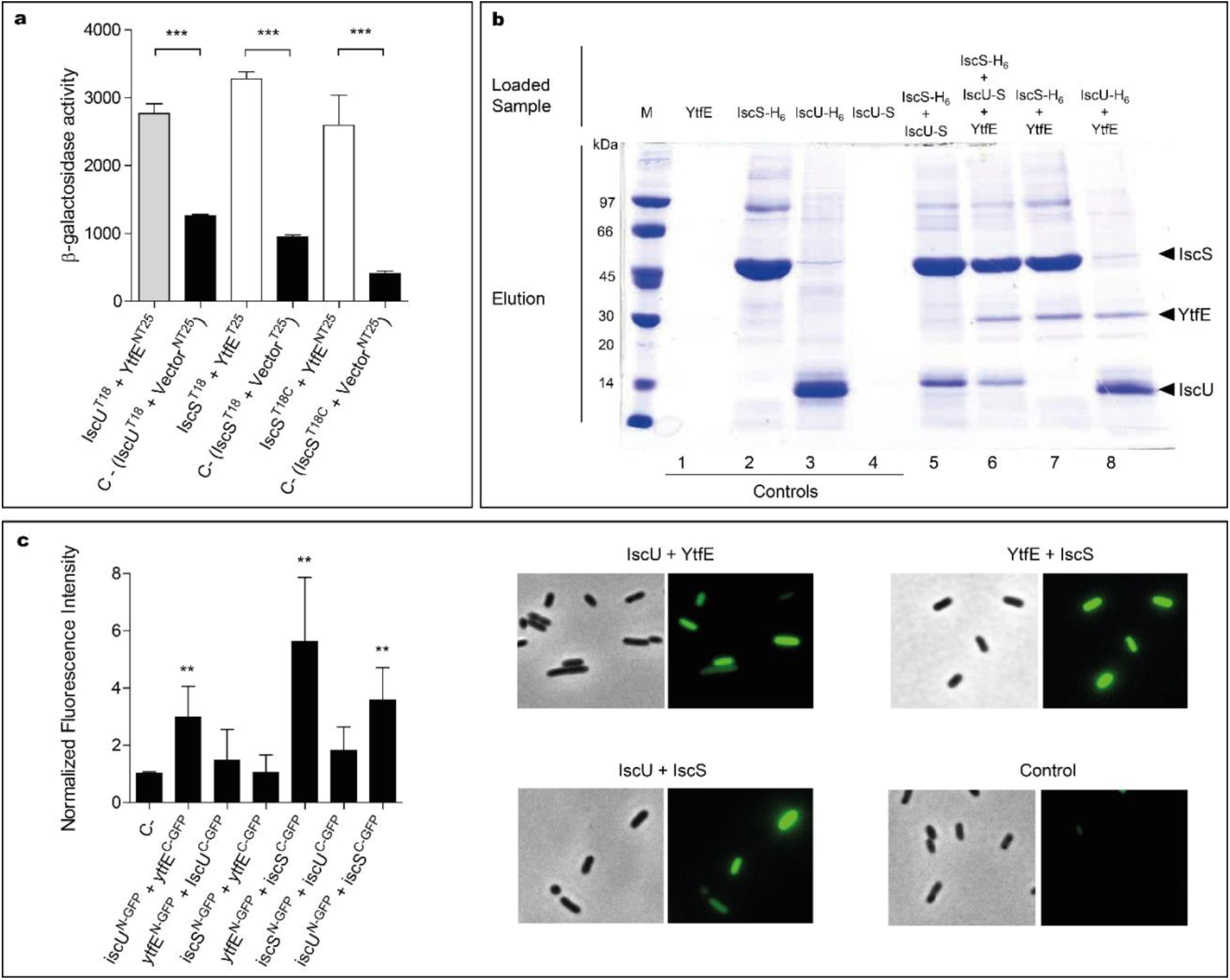
Analysis of the interaction of YtfE with IscS and IscU. **a**. The efficiency of the functional complementation between hybrid proteins was measured by β-galactosidase assays. The interaction of YtfE with IscU (grey bar) and IscS (white bars) was evaluated in *E. coli* cells co-transformed with plasmids containing either *iscU* or *iscS* genes cloned at the N- or C-terminal of the T18 fragment (T18/T18C) and the plasmid harbouring the *ytfE* gene cloned at the N- or C-terminal of the T25 fragment (NT25/T25). T25 empty plasmid together with the plasmid containing either the *iscU* or the *iscS* gene were utilized as negative controls (black bars). Values are means ± standard errors from at least three independent cultures analysed in duplicate. ***, P < 0.0001; **b**. For pulldown assays of YtfE, IscS and IscU, samples are obtained from cell extracts expressing one, two or three proteins loaded in Ni-chelating columns, eluted with 1 M imidazole buffer, and subjected to SDS-PAGE. Lanes 1-4: control samples from extracts expressing only one protein. Lanes 5-8: co-purification of IscS with IscU (5); co-purification of YtfE and IscU with IscS (6); co-purification of IscS with YtfE (7); and co-purification of IscU with YtfE (8); **c**. In BiCF of YtfE and ISC proteins, cells were co-transformed with vectors expressing YtfE and IscS or IscU – GFP fusions. On the left, fluorescence quantification was performed using MetaMorph Microscopy Automation and Image Analysis Software. Fluorescence values for the negative control (empty plasmid vectors) were normalized to 1. Values are means ± standard errors from at least three independent cultures analysed in duplicate. **, P < 0.005. On the right microscope images (Bright field phase and FITC) of Negative control; iscU^N-GFP^ + YtfE^C-GFP^; ytfE^N-GFP^ + IscS^C-GFP^; and iscU^N-GFP^ + IscS^C-GFP^.

A pulldown assay was also used to show the interaction between YtfE and Isc proteins. In this method, we grew cells transformed with plasmids expressing YtfE non-tagged, IscS and IscU (both fused at the N-terminal to a H6-tag) and IscU linked C-terminally to a S-tag. Extracts from cells expressing only one of the proteins were treated and analysed similarly to serve as controls. Cell extracts were loaded onto Ni-chelating columns and proteins were eluted at 1 M imidazole, and the collected fractions were analysed by SDS-PAGE (Fig. 1b).

Cells containing IscS-H6 or IscU-H6 strongly bind to the matrix and were eluted at 1 mM of imidazole (Fig. 1b, lanes 2-3). On the contrary, cells expressing, separately, YtfE or IscU-S do not bind to the resin (lanes Fig. 1b, 1 and 4). As expected, IscS-H6 retains IscU-S due to the interaction of the two proteins (Fig. 1b, lane 5), a result that is in agreement with previous studies that showed the IscU/IscS interaction by other methods(21–23).

We observed that both IscS-H6 and IscU-H6 retain the non-tagged YtfE (Fig. 1b, lanes 7 and 8, respectively). Furthermore, when cells express simultaneously IscS-H6, non-labelled-YtfE and IscU-S (in which the last proteins cannot bind by itself to the Ni-resin) co-elution of the three proteins was observed (Fig. 1b lane 6), further proving their interaction and suggesting the formation of a three-protein complex.

Lastly, we used Bimolecular Complementation Fluorescence (BiFC), that is based on the reconstitution of the GFP protein, to analyse the interaction of the proteins *in vivo*. GFP-fragment fusions were linked to the N- and C-terminal domains of YtfE, IscU and IscS so that all possible combinations were tested. *E. coli* cells containing plasmids co-expressing YtfE/IscU, YtfE/IscS and IscS/IscU fusions, prepared as described in Methods, were spread onto agarose slides and the microscopy images and fluorescence intensity values are shown in Fig. 1c.

As expected, cells expressing IscU^N-GFP^/IscS^C-GFP^ exhibited a positive interaction with a 3-fold higher fluorescence than the control sample. More important, also the *E. coli* cells expressing separately IscU^N-GFP^/YtfE^C-GFP^ and YtfE^N-GFP^/IscS^C-GFP^ exhibited higher fluorescence values by approximately 3-fold and 6-fold relative to the control, respectively (Fig. 1c). Moreover, the interactions are shown to be conformationally dependent as cells expressing YtfE^N-GFP^/IscU^C-GFP^ and IscS^N-GFP^/YtfE^C-GFP^ had no significant fluorescence.

Altogether, the results clearly show that YtfE interacts with IscS and IscU that are two major proteins of the *E. coli* ISC system.

### E159 and E125 modulate the iron release from *E. coli* YtfE

Based on the data above that provides a physiological meaning to the iron donor capacity of YtfE, we ought to identify the amino acid residues that modulate these properties by using site-directed mutant proteins and determine their iron binding parameters. Our hypothesis was that these residues would be among the highly conserved amino acid residues that are located in the regions that connect the protein to the solvent. Based on the YtfE crystal structure(12), two residues could fulfil these requisites, namely E125 and E159. Thus, the wild-type protein and five YtfE proteins in which these residues were replaced by the neutral amino acid leucine and by positively or negatively charged residues, such as asparagine and aspartic acid, were prepared (YtfE-E125L, YtfE-E125N, YtfE-E125D, YtfE-E159L, YtfE-E159N). Furthermore, to investigate the impact on the iron releasing properties of the N-terminal region of YtfE that is closely located to a possible second exit tunnel(12) (see below), the YtfE^Truncated^ and YtfE^M^ mutant proteins were also studied.

Proteins exhibited an iron content of ~2 and their UV–visible spectra contained a broad band approximately at 360 nm (data not shown), that exhibits features similar to those of the wild-type YtfE(2, 9). The only exception was the YtfE-E159L mutant that had a lower iron content, which is in agreement with our previous results(9).

To determine the iron binding chemistry, the proteins were incubated with the iron chelator desferrioxamine and the formation of the desferrioxamine-Fe(III) complex was monitored by following the intensity of the 420 nm band. The dissociation constant K_d_ of Ytfe-Fe(III) was calculated using the equation described in Methods. In addition, the kinetics of YtfE iron release was also determined through the initial ferric iron release rate (V_0_), which was calculated from the slope of a linear fit to the curve obtained when using the highest concentration of chelator (1000 μM) (Table 1, Supplemental Fig. S1 and S2).

**Table 1:**
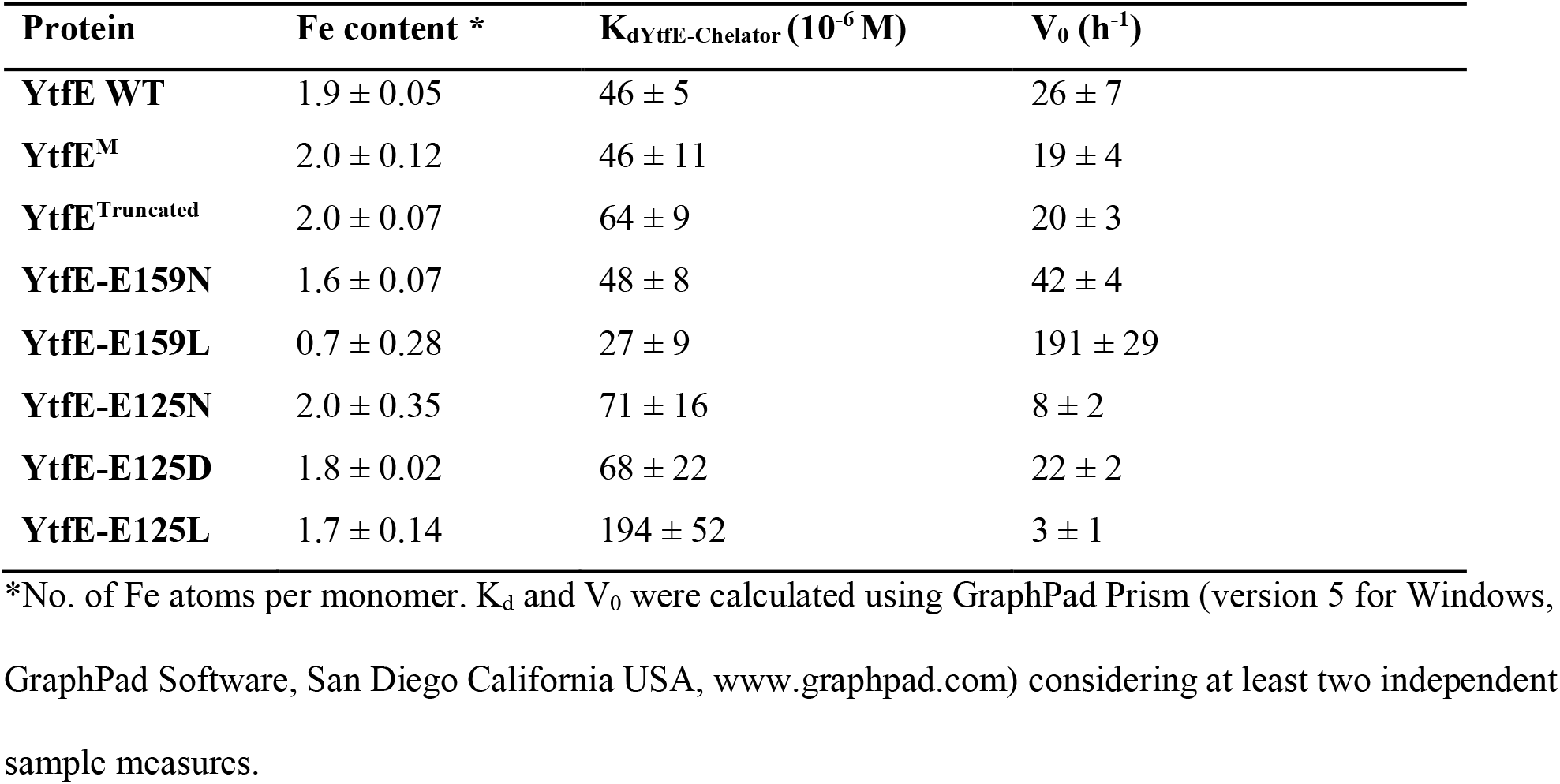
Iron content, dissociation constants and initial iron release rates of *E. coli* YtfE and variants.

| Protein | Fe content * | K<sub>d</sub><sup>YtfE-Chelator</sup> (10<sup>-6</sup> M) | V<sub>0</sub> (h<sup>-1</sup>) |
| --- | --- | --- | --- |
| **YtfE WT** | 1.9 ± 0.05 | 46 ± 5 | 26 ± 7 |
| **YtfE<sup>M</sup>** | 2.0 ± 0.12 | 46 ± 11 | 19 ± 4 |
| **YtfE<sup>Truncated</sup>** | 2.0 ± 0.07 | 64 ± 9 | 20 ± 3 |
| **YtfE-E159N** | 1.6 ± 0.07 | 48 ± 8 | 42 ± 4 |
| **YtfE-E159L** | 0.7 ± 0.28 | 27 ± 9 | 191 ± 29 |
| **YtfE-E125N** | 2.0 ± 0.35 | 71 ± 16 | 8 ± 2 |
| **YtfE-E125D** | 1.8 ± 0.02 | 68 ± 22 | 22 ± 2 |
| **YtfE-E125L** | 1.7 ± 0.14 | 194 ± 52 | 3 ± 1 |

\*No. of Fe atoms per monomer. K<sub>d</sub> and V<sub>0</sub> were calculated using GraphPad Prism (version 5 for Windows, GraphPad Software, San Diego California USA, www.graphpad.com) considering at least two independent sample measures.

Table 1 shows that the K_d_ values of YtfE, YtfE^M^ and YtfE-E159N do not differ substantially, while YtfE-E159L presents a slightly lower value that indicates a more loosely bound iron. On the contrary, YtfE^Truncated^ and the proteins containing mutations in E125 exhibited higher K_d_ values. In particular, the ferric dissociation constant of YtfE is substantially modified by the E125L mutation, becoming about 4-fold larger than that determined for the wild-type protein. Therefore, E125 greatly influences how strongly Fe(III) is bound to the YtfE scaffold.

Results in Table 1 also show that the kinetics of Fe(III) release from YtfE is controlled by E159 and E125, and is greatly influenced by the nature of the substituent residue. Glutamate is a polar charged residue, and its replacement by the polar but uncharged residue asparagine increased the iron release rate; an even higher increase was observed (by *ca*. 7-fold) when E159 was substituted by the hydrophobic residue leucine.

Interestingly, an opposite effect occurred upon substitution of E125, originating proteins with a slower iron release rate. Mutation of E125 to asparagine decreased the V_0_ values by about 3-fold whereas a leucine substitution reduced the iron-release rate by *ca*. 8-fold.

Altogether, these results revealed that glutamate E125 is a key role in the modulation of the thermodynamic and kinetic properties of the iron release from the YtfE di-iron centre.

### E125 is essential for YtfE to act as iron donor

Considering the results described above, we determined the impact of the E125 mutation on the YtfE iron donor capacity to assist formation of a Fe-S centre in the ISC system. With this in mind, reactions mixtures containing *E. coli* apo-IscU, YtfE (wild-type or mutant proteins), IscS, L-cysteine and DTT (1,4-dithiothreitol), were monitored under anaerobic conditions and the cluster formation in IscU was monitored by visible spectroscopy.

The spectrum of the reaction mixture containing IscU and wild-type YtfE, obtained after a period of incubation of 1 h, exhibited bands at ~456 nm and ~410 nm characteristic of the presence of a [2Fe-2S]^2+/1+^cluster (Supplemental Fig. S3). On the contrary, the spectra of the reaction mixtures containing IscU and YtfE-E125N or IscU and YtfE-E125L exhibited very weak bands (Supplemental Fig. S3), which indicates that the amount of iron-sulfur cluster formed in IscU is negligible.

These results are consistent with the thermodynamic and kinetic properties of YtfE-E125N and YtfE-E125L which show that in these proteins the iron is more tightly bound and the rate of iron release is much lower.

Hence, the mutation of E125 by a hydrophobic or a positively charged residue hindered the iron donation capacity of YtfE which led us to conclude that residue E125 plays a key role on the protein function.

### Structural basis for the iron donor properties of YtfE

The unexpected iron binding properties of the YtfE-E159 and YtfE-E125 mutant proteins, in which the modified amino acid residues are located outside of the coordination sphere of the di-iron centre (Fig. 2a), prompted us to address the structural role of two glutamates in YtfE. We resorted to X-ray crystallography to determine their structures and, for comparison purposes, we revisited the structure of the wild-type YtfE.

**Fig. 2.**
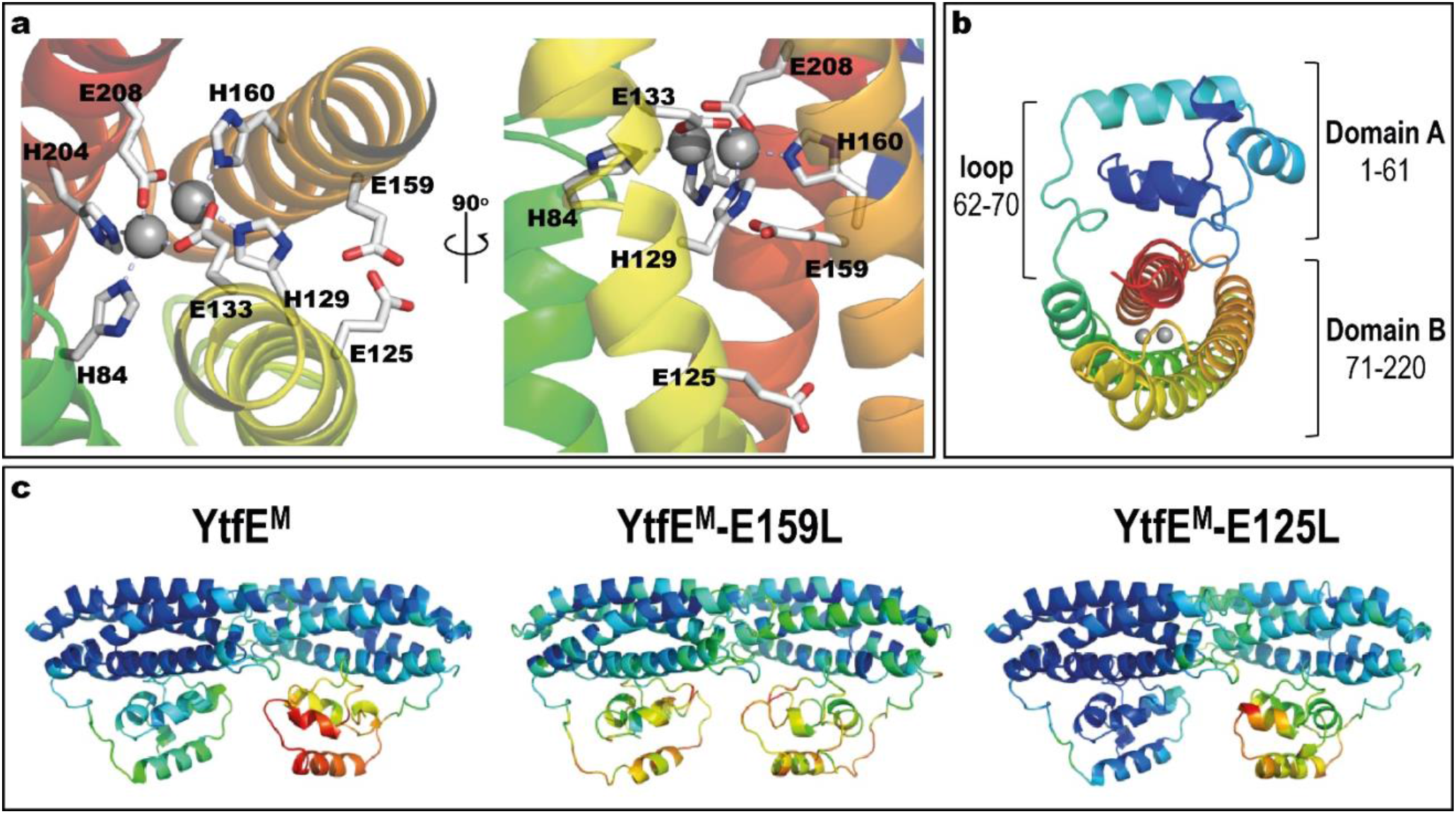
*E. coli* YtfE^M^ overall structure. **a.** Detail views of the YtfE^M^ di-iron centre, its coordinating residues, and residues E125 and E159 are highlighted in light grey (left panel view is rotated 90 ° about a vertical axis from first panel). **b**, Cartoon representation of the *E. coli* YtfE^M^ monomer (chain A) showing the two domains that are connected by a loop. In **a** and **b**, the monomer structure is rainbow coloured from dark blue in the N-terminal to red in the C-terminal. The di-iron site is represented as grey spheres. **c**, Cartoon representation of the *E. coli* YtfE^M^, YtfE^M^-E159L and YtfE^M^-E125L subunits present in the asymmetric unit, coloured according to their B-factor values (the di-iron site is omitted).

Crystals of *E. coli* YtfE could only be obtained when the proteins contained the mutation to alanine of the two vicinal cysteines C30 and C31 (herein designated YtfE^M^), a result that agrees with previous reports(12). Therefore, we prepared crystals of YtfE^M^, YtfE^M^-E159L and YtfE^M^-E125L as described in Methods. The atomic coordinates and experimental structure factors were deposited in the Worldwide Protein Data Bank(24) with the accession codes 7BHA, 7BHB and 7BHC for the YtfE^M^, YtfE^M^-E159L and YtfE^M^-E125L structures, respectively. Data collection and refinement statistics are listed in Supplemental Tables 1 and 2.

The three crystal structures contain two molecules in the asymmetric unit (chains A and B) in monoclinic space group *P*2_1_. All molecules are composed of two domains (A and B) and have continuous electron density for the full length of the protein (220 residues) (Fig. 2b). However, in all the structures here presented, the electron density for chain B is overall weaker than for chain A, as inferred from their mean B-values (see Supplemental Table 3) and therefore only chain A will be considered for the following structural analysis. The crystal structure of YtfE^M^-E125L resembles more that of YtfE^M^. However, in YtfE^M^-E159L more regions with low density are observed, and the highest B-values are associated with domain A (Fig. 2c, Supplemental Table 3).

Domain A, usually referred to as a ScdA_N-like domain(25, 26) is composed of 4 α-helices, and contains the DxCCG motif in the loop between helices 2 and 3, which is highly conserved in the RIC family (Supplemental Fig. S4). The structures exhibited a domain A fold similar to the previous reported YtfE wild-type structure(12) with a r.m.s.d. (root-mean-square deviation) of only 0.3 Å between 188-193 superposed main-chain Cα atoms.

In the three structures, several residues in domain A can establish intermolecular contacts with symmetry-related monomers. In YtfE^M^, hydrogen bonds occur between residues D5 and R17, E49 and R86. In the YtfE^M^-E125L structure, the same hydrogen bonds are present together with bonds between R39 and K57/E60/Q61. Interestingly, in the YtfE^M^-E159L structure, no such hydrogen bond interactions are present, probably explaining why the B-values in this domain are higher than in the other structures (Fig. 2c, Supplemental Table 3).

Like for domain A, the overall four-helix bundle structure of domain B is maintained independently of the mutation introduced in the protein. As noted earlier(9, 10, 27), domain B shares a similar structural topology with proteins such as hemerythrins, rubrerythrins and (bacterio)ferritins, with a r.m.s.d. of *ca*. 3 Å between superposed main-chain Cα atoms (Supplemental Fig. S5). An extensive structural analysis of the available hemerythrin-like proteins(27), rubrerythrins, and (bacterio)ferritins, revealed that the arrangement of the helices in YtfE is unique in its left-handed four-helix bundle structure. Hemerythrins have right-handed four-helix bundles (Supplemental Fig.S5b), and rubrerythrins or (bacterio)ferritins exhibit a mixed arrangement, starting as left-handed and then becoming a right-handed four-helix bundle (Supplemental Fig. S5c,d). These results show that in relation to the structural fold of the four-helix bundle, YtfE and RIC proteins form a separate cluster within the hemerythrin and ferritin-like families.

Analysis of the electrostatic potential on the molecular surface of *E. coli* YtfE^M^ shows that domain A is mainly positively charged and domain B is mainly negatively charged (Fig. 3a,b). In addition, the cavity where the di-iron centre is inserted is positively charged and the negative charge at the surface in this region is mainly due to residues E125, E159 and E162 (Fig. 3c).

**Fig. 3.**
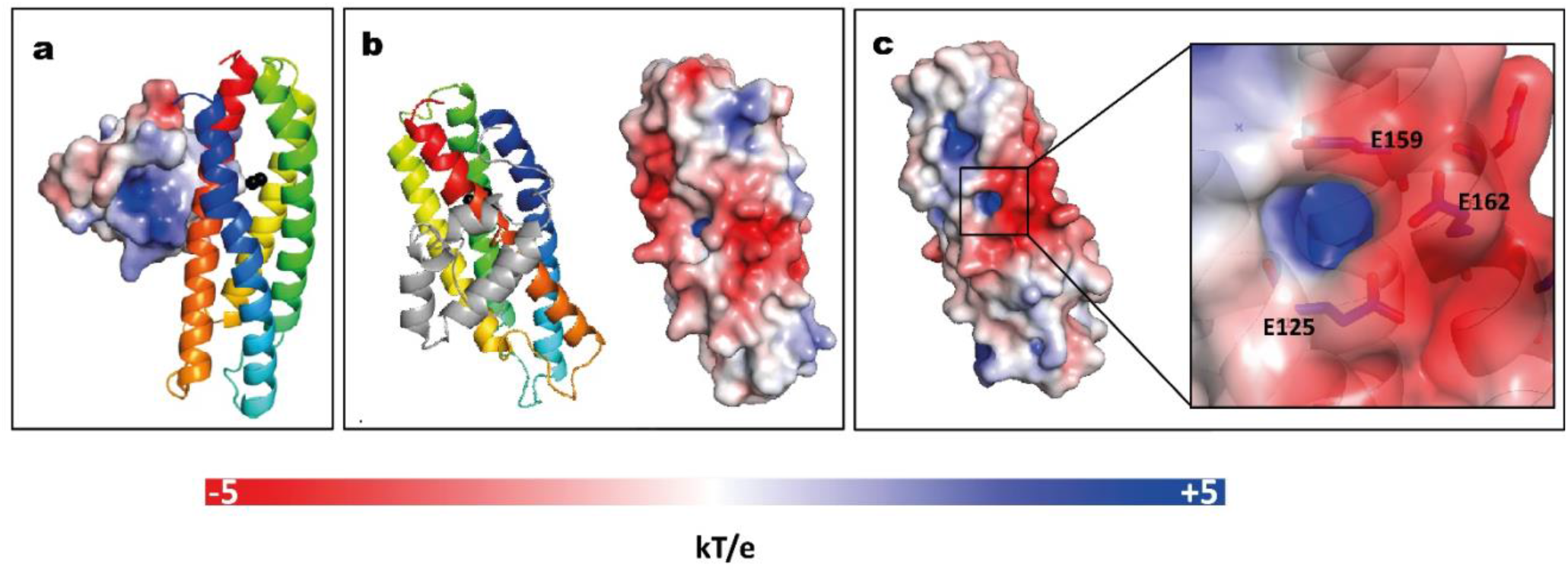
Electrostatic potential at the molecular surface of *E. coli* YtfE^M^. **a**, Electrostatic potential surface of domain A that faces domain B is positively charged. **b**, View of YtfE^M^ rotated 90° about a vertical axis from panel **a**, the electrostatic surface representation (right) is on the same orientation as the cartoon representation of the structure (left), without domain A. **c**, Detail of the electrostatic surface (negatively charged) near the iron cavity (positively charged). The electrostatic potential at the molecular surface was obtained considering that YtfE^M^ is loaded with Fe^3+^-Fe^2+^. The electrostatic potential surfaces range from −5 kT/e (red) to + 5 kT/e (blue). Figure prepared with PyMOL(56) using ABPS(58) and PDB2PQR(59) server to generate the electrostatic potential surface. The charge for each metal was manually added to the PQR file, taking in consideration the oxidation state of the protein.

### Structural features of the di-metal site

*E. coli* YtfE^M^ and YtfE^M^-E125L contain a di-iron site coordinated by four histidine residues, through their N^ε2^ atoms (H84 and H204 for Fe1, H129 and H160 for Fe2), and two bidentate glutamic acids that bridge the two irons (E133, E208) (Fig. 4a,c). These residues are part of the H84(x_44_)H129(x_3_)E133(x_26_)H160(x_43_)H204(x_3_)E208 binding motif, that shares similarities with the four-histidine binding motif in hemerythins. However, in YtfE the two bridging ligands are glutamates, whereas in hemerythrins one residue is a glutamate and the other is an aspartate (27). Besides these coordinating ligands, an oxo-bridge between the iron atoms is also present in YtfE. Except for YtfE^M^-E159L that only contains one iron atom in the centre, the di-iron coordination and the distances of the ligands to the irons are similar in the YtfE^M^ and YtfE^M^-E125L structures. Supplemental Table 4 lists details of the coordinating distances of the structures.

**Fig. 4.**
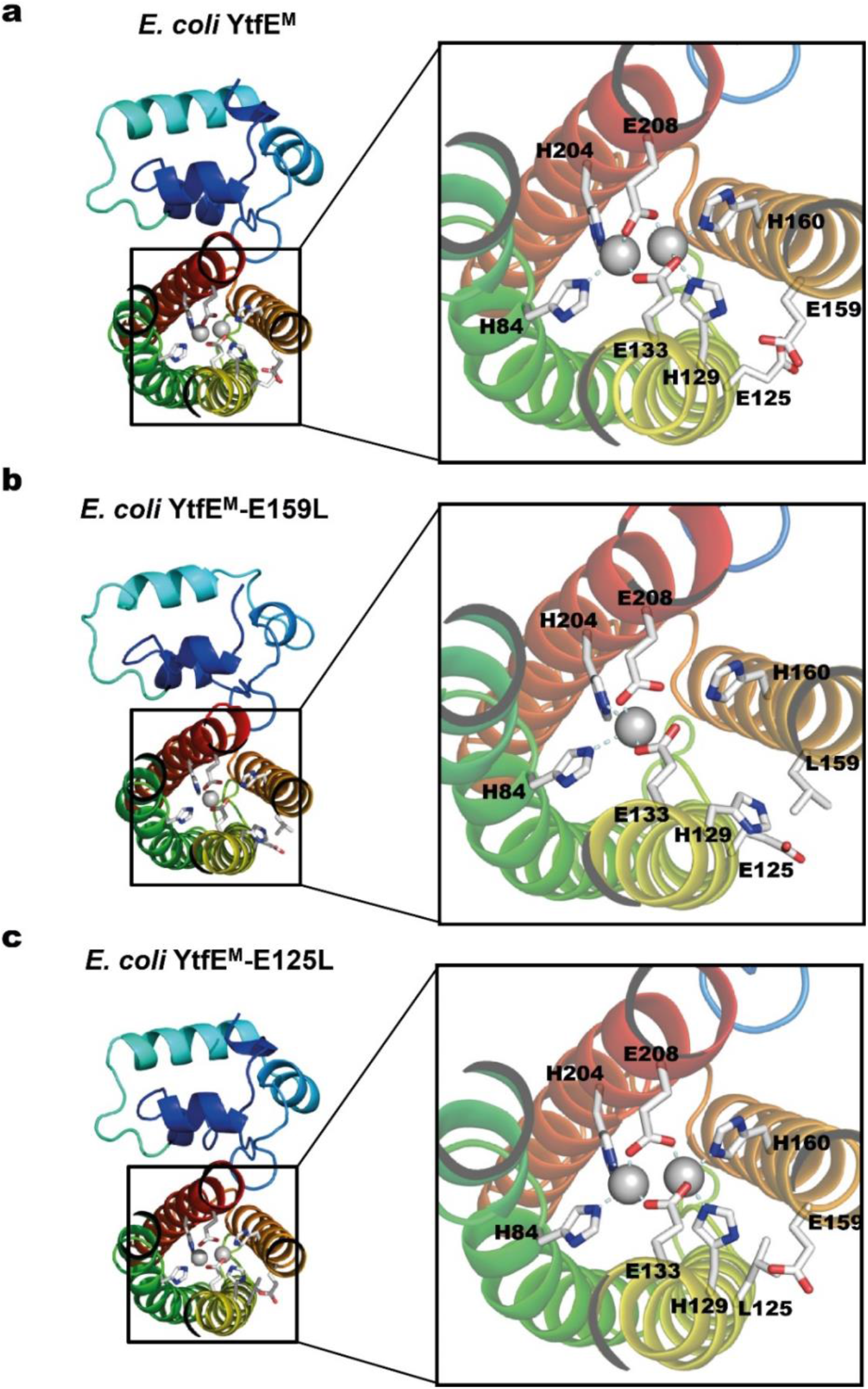
*E. coli* YtfE Iron centre. **a**, YtfE^M^ di-iron centre, coordinated by residues H84, H129, E133, H160, H204 and E208. **b**, YtfE^M^-E159L mononuclear Fe centre coordinated by H84, H204 and E208. The second Fe atom is not present and residue H129 assumes a different side chain conformation from that in YtfE^M^. **c**, In YtfE^M^-E125L the di-iron centre, is similar organized as in YtfE^M^.

As mentioned above, YtfE^M^-E159L contains only one iron atom in the metal site (Fig. 4b, Supplemental Fig.S6). Additionally, the conformation of ligand H129 differs from that observed in the other two proteins. Specifically, the side chain of H129 is ~ 7 Å apart from its original position and no longer pointing towards the di-iron centre (the closest distance to YtfE^M^-E159L H129 is *ca*. 9 Å whereas in Ytfe^M^ it is about 2 Å).

### The iron exit channel is controlled by E125

The structure of *E. coli* YtfE^M^ contains two channels(12). One channel, mainly hydrophobic (which will not be addressed in this study), and a hydrophilic channel that connects the metal centre to the solvent (Fig. 5a). The di-iron centre is located *ca*. 10 Å below the channel entrance, which is approximately circular in shape with a diameter of ~2.3 Å. The channel is formed mainly by hydrophilic residues, namely H160 and H129 (that are also Fe ligands), K132, E159, E125 and E162. The three negatively charged glutamates are located at the channel entrance and solvent exposed suggesting a possible iron exit(12). Hence, mutation of these residues was done and the channel in YtfE^M^ was compared with that present in the YtfE^M^-E159L and YtfE^M^-E125L mutant proteins.

**Fig. 5.**
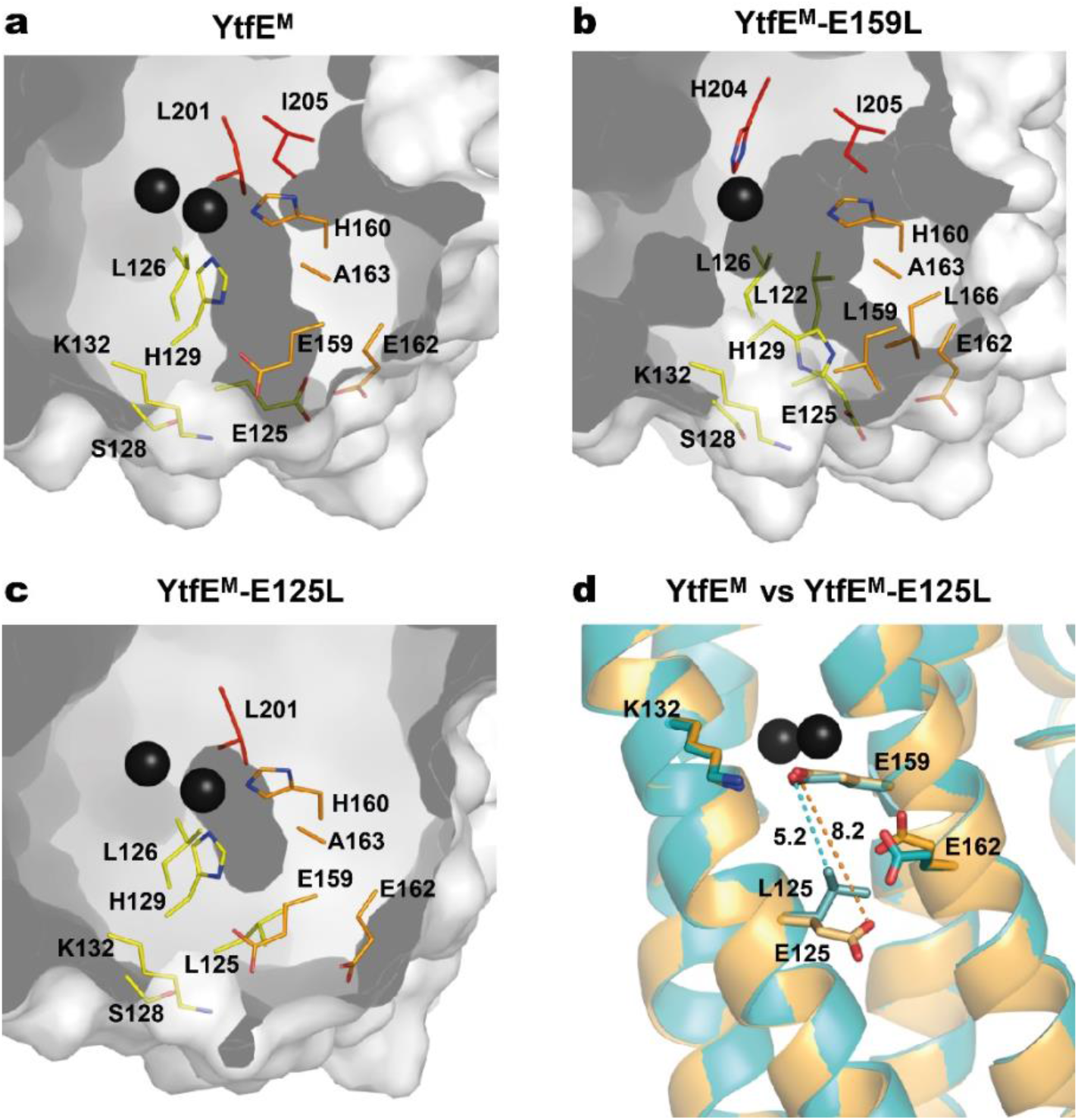
Channels present in YtfE^M^ and YtfE^M^-E159L structures. Channels are represented in dark grey. Residues lining the iron channel are represented as sticks with carbon atoms rainbow coloured according to the helices (helix 2 - yellow, helix 3 - orange and helix 4 - red). **a**, YtfE^M^ iron channel connects the external surface to the di-iron site. YtfE^M^ has its channel entrance formed by residues E159, E162, E125 and K132. **b**, In YtfE^M^-E159L, the channel connects the external surface to the mononuclear iron centre, and has its entrance between residues E162, L166 and E125, i.e., slightly below that of the YtfE^M^ pore. **c**, No channels were detected in the YtfE^M^-E125L structure. A small cavity is observed near the di-iron centre, however, the channel is interrupted by mutation of glutamate 125 to leucine. **d**, Superposition of the YtfE^M^ (orange) and YtfE^M^-E125L (cyan) structures showing the distances between the residues involved in the channel entrance. Distance between YtfE^M^ residues 159 and 125 is represented by a dashed orange line whereas the distance between YtfE^M^-E125L residues is indicated by a dashed cyan line. The channels were modelled using MOLE(57) and PyMOL(56) software.

The structure of YtfE^M^-E159L mutant contains only one iron atom in the metal centre that is located *ca*.13 Å below the channel entrance. In this case, the channel is longer, with a length of about 16 Å, and the channel entrance although retains the circular shape its diameter is reduced to ~ 1.5 Å (Fig. 5b). The channel is also formed by hydrophilic residues, namely by the ligands H204, H129, H160, and residues E125, E162 and L166 at the channel entrance. In YtfE^M^-E159L, the absence of the second Fe atom, the different position of the H129 sidechain (mentioned above) and the presence of leucine in position 159 modified the channel arrangement so that the substituting L159 residue is no longer included in the channel (Fig. 5a,b and Supplemental Fig.S6).

The mutation YtfE^M^-E125L induced a major modification of the channel (Fig. 5c). In this case, a significant shortening of the distance between the sidechains of residues L125 and E159 is observed, from 8.5 Å in the YtfE^M^ structure to 5.3 Å in the YtfE^M^-E125L mutant structure (Fig. 5d). As a consequence, the new position of L125 results in the occlusion of the channel and, consequently, the di-iron centre is no longer exposed to the solvent. The closing of the channel is consistent with the different iron release properties of the YtfE^M^-E125L, and explains why this mutant protein is no longer able to promote the formation of the Fe-S centre in the IscS, as described above.

## Discussion

Previously, we showed that *E. coli* YtfE is a major candidate to be an iron donor for IscU, based on two experimental observations. First, the YtfE dissociation constant for iron is lower than those of other potential iron donors, namely CyaA, YggX, IscA and SufA (11, 28–31). Second, YtfE can provide iron to IscU in Fe–S cluster reconstitution assays via a cysteine-mediated process. In this work, we proved that YtfE interacts with IscU and IscS, and revealed the structural features that underpin the YtfE iron donation properties.

We reanalysed the wild-type YtfE^M^ and obtained the YtfE^M^-E125L and YtfE^M^-E159L structures. Although the mutations did not change the overall structure, the single replacement by uncharged amino acids of the negatively charged amino acid residues E159 and E125, which are not ligands of the di-iron centre, introduced significant changes locally which were sufficient to impair the function of the protein. In *E. coli* YtfE, the highly conserved residue E159 is located at about 7 Å from the closest iron atom, therefore it is not involved in metal coordination. Replacement of E159 with another polar amino acid, YtfE-E159N, generated a protein with K_d_ values similar to those of the wild-type. However, in YtfE-E159L with a neutral leucine instead of the glutamate, the metal site lost one of its iron ions. Thermodynamic and kinetic data indicate that mutation of E159 generates a protein with a more unstable centre that releases iron at a higher rate. Moreover, the structure of YtfE^M^ revealed that when E159, that establishes a hydrogen bond with the iron ligand H129 (H129^Nδ1…^E159^Oε1^-2.66 Å), is replaced by leucine, the position of the neighbouring histidine H129 alters its sidechain conformation, pointing away from the centre and towards the protein surface (Fig. 4b, Supplemental Fig. S6)). Thus, the intermolecular interaction between H129 and E159 is crucial for the overall stability of the centre. It is also interesting to note that in our previous work, H129 appeared to be the only histidine ligand that controls the ability of YtfE to donate iron(9).

The mutation of E125 (which is located at *ca*. 9 Å from its nearest iron in the di-iron centre) to leucine generated a protein with an intact binuclear site. However, and unexpectedly, the YtfE-E125L mutant has a hindered capacity to release iron and is not able to promote the assembly of the iron-sulfur centre in IscU. These results were rationalized by the analysis of crystal structure of YtfE^M^-E125L showing that the positioning of the L125 sidechain blocks the external part of the hydrophilic channel, and consequently impairs the release of iron from the di-iron cluster to protein exterior.

Thus, we have identified a mechanism through which a di-iron protein has the possibility of releasing iron as long as it contains in the upper part a set of negatively charged glutamate residues forming a channel that links the di-iron centre to the surface. The mononuclear iron YtfE-E159L mimics an YtfE protein following iron release and its structure shows that the process implies the movement of one of the axial histidines that exposes the metal site cavity to the exterior, thus making possible the re-entry of another iron atom to rebuild the di-iron centre. Since we previously showed that YtfE interacts with *E. coli* Dps (DNA-binding Protein from Starved cells) (32), we may hypothesise that this iron storage protein constitutes a good candidate to promote the reconstitution of the di-iron cluster in YtfE.

An identical mechanism is proposed to occur in the proteins of the RIC family, as residues equivalently located to H129, E125 and E159 are conserved in the vast majority of RICs. Furthermore, our results open the possibility that this a common dynamic behaviour of di-iron proteins that share similar structural features, such as bacterioferritins(33). For example, when *Desulfovibrio desulfuricans* bacterioferritin is fully reduced and subsequently allowed to oxidize in the presence of atmospheric oxygen, it loses one of the iron atoms of the ferroxidase centre(34) through a not yet identified mechanism. Like RIC proteins, (bacterio)ferritins also contain a pore 6 to 7 Å deep that connects directly the di-iron ferroxidase centre to the surface, that has been proposed to be involved in the entry of ferrous iron to the centre(34–38). Moreover, the movement of histidine and glutamic acid residues is also observed and predicted to be related to the opening of the ferroxidase centre to the solvent(34–36). Our observations for YtfE could explain how bacterioferritins might donate iron from their di-iron centre.

Despite the importance in all organisms of the Fe-S assembly systems and, in particular, of the ISC system operative in bacteria, the nature of the iron donor has remained elusive. Although frataxin (CyaA) was initially proposed to function as an iron storage/donor for IscU, the most recent studies indicate that frataxin is mainly an accelerator of persufilde transfer(39). Other candidates for iron donors belong to the A-type protein family that includes IscA, NifA and SufA from various organisms, which were shown to bind ferric iron with high affinity and to provide iron to IscU. However, the IscA proteins interact only with the late-acting components of the Fe–S cluster biogenesis pathway, which has reinforced the idea that they are not involved in iron insertion(40, 41). In this work, we prove that YtfE interacts with IscU and IscS, and we have identified in the structure of YtfE a channel through which the iron may exit to promote the formation of the cluster assisted by Isc proteins. Altogether, the results now presented strongly support the proposal that in a large number of bacteria the di-iron RICs act as iron donors for the biogenesis of Fe-S clusters.

## Methods

### Protein-protein interaction experiments

#### Bacterial two-hybrid (BACTH) system

The system based on the interaction-mediated reconstitution of the *Bordetella pertussis* adenylate cyclase (Cya) activity in *E. coli*(20) was used to analyse the interaction of YtfE with IscU and IscS proteins.

The genes coding for IscS, IscU, and YtfE were PCR-amplified using appropriate pairs of oligonucleotides (Supplemental Table 5) and genomic DNA from *E. coli* K12 as templates. The generated DNA fragments were digested with BamHI and KpnI (for the *iscS* and *iscU genes*) or with BamHI and SalI (for the *ytfE gene*), and cloned into the corresponding sites of pKT25, pKNT25, pUT18 and pUT18C (Supplemental Table 6). The recombinant plasmids allowed the production of proteins linked to the C- and N-termini of the T25 domain and to the N- and C-termini of the T18 domain of *B. pertussis* Cya. All recombinant clones were sequenced confirming the absence of undesired mutations. The complementary plasmids (carrying a T25 fragment and a T18 fragment in all possible combinations) were co-transformed into *E. coli* DHM1, a strain deleted in the *cya* gene. The pUT18 and pKT25 empty plasmids co-transformed with the complementary plasmids containing *iscS* or *iscU* were used as negative controls.

The efficiency of interactions between the hybrid proteins was quantified by the β-galactosidase activity method using *E. coli* DHM1 as the recipient strain. All β-galactosidase assays were done in LB broth supplemented with antibiotics. At least 3-4 representative colonies of each transformation plate were inoculated, in duplicate, in LB broth and, following an overnight growth at 37 °C, cells were re-inoculated 1/100 into LB containing ampicillin, kanamycin and isopropyl-1-thio-β-D-galactopyranoside (IPTG). When cells reached an OD_600_~0.5 (approximately after 16 h growth at 30 °C), 1 mL of each culture was collected by centrifugation (5 min, 5000 rpm). The pellets were suspended in 100 μL BugBuster HT 1x (Novagen) for cellular lysis and incubated at 37 °C for 30 min. Cellular debris were removed by centrifugation and each suspension (20 μL) was tested in duplicate for the enzymatic reaction in a microtiter plate reader. The β-galactosidase assays were initiated upon addition of the following mixture: 0.27%β-mercaptoethanol (v/v), 0.9 mg/mL ONPG (o-nitrophenyl-β-D-galactopyranoside) in buffer Z (60 mM Na_2_HPO_4_, 40 mM NaH_2_PO_4_, 1 mM MgSO_4_ and 10 mM KCl). The absorbance was measured at 420 nm every 2 minutes, and the reaction continued at 28 °C for 1.5 h. The β-galactosidase specific activity is defined in units per milligram of protein. For the conversion of microplate reader Abs420 values into standard spectrophotometer values, a correction factor of 2.2 was determined using serial dilutions of an O-nitrophenyl (ONP) solution. Positive interactions were considered for β-galactosidase activities at least two times higher relative to the negative control.

#### Pulldown assays

The genes encoding YtfE, IscS and IscU were amplified from *E. coli* K-12 genomic DNA by PCR using specific oligonucleotides, cloned into pFLAG-CTC (Sigma), and in pACYCDuet-1 (Novagen) that allows co-expression of two target genes fused with S-tag (S) and His_6_-tag (H6) sequences (Supplemental Tables 5,6). Sequencing of the recombinant plasmids confirmed their integrity and the absence of undesired modifications. *E. coli* BL21(DE3) Gold (Agilent) was transformed with one or two plasmids: (i) pFLAG-ytfE (expressing YtfE); (ii) pACYC-IscS-H6 (expressing IscS fused to a N-terminal H6-tag); (iii) pACYC-IscU-H6 (expressing IscU fused to a N-terminal H6-tag); (iv) pACYC-IscU-S (expressing IscU fused to a C-terminal S-tag); (v) pACYC-IscS-H6-IscU-S (expressing IscS with a H6-tag N-terminal and IscU with a C-terminal S-tag); (vi) pFLAG-ytfE and pACYC-IscS-H6; (vii) pFLAG-ytfE and pACYC-IscU-H6; and (vii) pFLAG-ytfE and pACYC-IscS-H6-IscU-S. Cells harbouring single recombinant plasmids (from i to iv) were used as control samples. Growth was done at 30 °C in Terrific Broth (TB) medium supplemented with antibiotics and 100 μg/mL of Fe citrate. When the culture reached an OD_600_ of 0.3-0.4, 0.3 mM IPTG were added to induce the expression of the YtfE, IscS and IscU proteins. After 5 h, the cross-linking agent formaldehyde (1% final concentration) was added. Incubation proceeded at 37 °C for 20 min, and the reaction was stopped by incubation with 0.5 M glycine at room temperature for 5 min(32). Cells were harvested by centrifugation, washed twice, resuspended in PBS, disrupted in a French press (Thermo Scientific), and debris removed by centrifugation (15000 rpm for 30 min). The total protein concentration of the supernatants was determined by the Pierce bicinchoninic acid (BCA) protein assay (Thermo Scientific). For the pulldown experiments, the supernatants were loaded into Ni-chelating Sepharose Fast Flow columns (GE Healthcare), which were previously equilibrated with 20 mM Tris-HCl (pH 7.9) supplemented with 0.5 M NaCl (buffer A) containing 5 mM imidazole. Columns were washed with buffer A with 60 mM imidazole, and the proteins were eluted in the same buffer with 1 M imidazole(21). The protein fractions were analysed by SDS-PAGE (12.5%).

#### Bimolecular Complementation Fluorescence Assays (BiCF)

BiFC assays were done essentially as previously described(32, 42). Briefly, genes encoding YtfE, IscS and IscU were PCR-amplified from genomic DNA of *E. coli* K-12 using the oligonucleotides described in Supplemental Table 5. DNA fragments were cloned into vectors pET11a-link-N-GFP (using XhoI/BamHI sites) and pMRBAD-link-C-GFP (using NcoI/AatII sites)(42), which express the green fluorescent protein (GFP), to construct GFP fusion proteins located at the N- or C-termini, respectively. All recombinant plasmids were sequenced to confirm the integrity of the genes and the absence of undesired mutations. *E. coli* BL21(DE3) Gold cells (Agilent) were co-transformed with recombinant pET11a-N-GFP and pMRBAD-C-GFP vectors so that several combinations were tested, namely YtfE/IscS, YtfE/IscU and IscS/IscU. Colonies were inoculated in LB medium, grown overnight at 37 °C and 150 rpm, and plated on inducing and selective LB agar medium, which contained 20 μM IPTG, 0.2% of arabinose and adequate antibiotics. The plates were incubated overnight at 30 °C, followed by 2 days of incubation at room temperature. Colonies were suspended in phosphate-buffered saline (PBS) and spread onto 1.7% agarose slides. Samples were analysed for green fluorescence in a Leica DM6000 B upright microscope coupled to an Andor iXon+ camera, using 1000x amplification and a fluorescein isothiocyanate (FITC) filter. The images were analysed using the MetaMorph Microscopy Automation and Image Analysis Software (Molecular Devices).

### Site-directed mutagenesis of *E. coli* YtfE and production of proteins

Site-directed mutants of *E. coli* YtfE were constructed on the template pET-YtfE(2) using the oligonucleotide pairs shown in Supplemental Table 5, and the QuikChange II Site-Directed Mutagenesis Kit and protocol (Agilent Technologies). Reaction products were transformed into *E. coli* XL1-Blue competent cells (Agilent Technologies), and positive recombinant vectors were selected on agar-plates containing 30 μg/mL kanamycin. Primers were designed so that the YtfE codons for E159 and E125 were replaced by those of leucine, asparagine and aspartate. Also, cysteines C30 and C31 of YtfE were changed to alanines. The *E. coli ytfE* gene encoding the C30AC31A mutagenized protein (designated as YtfE^M^) was used to generate other mutants (YtfE^M^-E125L and YtfE^M^-E159L). Also, YtfE genes in which the codons for E159 and E125 were replaced by those of leucine, asparagine and aspartate were constructed. All plasmids were sequenced confirming the presence of the desired mutations and the absence of unwanted modifications. Table 1 summarizes the site-directed mutants of YtfE and YtfE^M^ proteins constructed in this work.

YtfE wild-type and its mutant proteins were expressed in *E. coli* BL21(DE3) Gold (Agilent) by growing cells in M9 minimal medium supplemented with 20 mM glucose and 0.1 μM FeSO_4_, under aerobic conditions and at 30 °C, to an OD_600_ of 0.3. At this point, 200 μM IPTG was added and expression was induced for 7 h. Cells were disrupted in a French Press, centrifuged and the soluble extracts loaded sequentially on a Q-Sepharose High-Performance and a Superdex S-75 gel filtration columns, coupled to an AKTA Purifier 10 FPLC System (GE Healthcare) following the protocol previously described(2). Proteins were judged pure by SDS-PAGE, and the iron content was determined by the 2,4,6-tripyridyl-S-triazine (TPTZ) method(43).

### Iron binding assays

To determine the Fe(III) dissociation constant and the rate of iron release from YtfE wild-type and site-directed mutants, the as-isolated proteins (25 μM) were incubated under aerobic conditions with the ferric chelator desferrioxamine (5–1000 mM, Sigma)(11). Reaction mixtures were prepared in 20 mM Tris-HCl pH 7.5 buffer with 150 mM NaCl, incubated at room temperature for 24 h to attain thermodynamic equilibrium and analysed by UV-visible spectroscopy in a Shimadzu UV-1700 spectrophotometer.

The K_d,YtfE-chelator_, defined by K_d,YtfE-chelator_ = [YtfE-Fe][Chelator]/([Chelator-Fe][YtfE]), is the dissociation constant of iron from YtfE in the presence of the competing chelator(11). The constant was determined with GraphPad Prism version 5 for Windows (GraphPad Software, San Diego California USA, www.graphpad.com) using the equation for a saturation binding experiment with one specific binding site: Y = B_max_^*^X/(K_d,YtfE-chelator_ + X), where X is the concentration of the chelator, Y the percentage of iron atoms released and Bmax the maximum percentage of iron atoms released in relation to the total number of iron atoms in the isolated protein. The extinction coefficient used for desferrioxamine-Fe(III) was 2.9 × 10^3^ M^-1^cm^-1^.The initial rate values (V_0_) for iron dissociation were calculated from the linear part of the curves fit to the experimental results, considering only the first hours of the reaction.

### Assembly of Fe-S clusters in IscU promoted by YtfE

*E. coli* M15:pREP4 cells (Quiagen) harbouring plasmids pQE30-(His)6-IscS and pQE60-IscU-(His)6 were used to produce *E. coli* IscS and apo-IscU proteins with a His-tag at the N- and C-terminal, respectively, and were purified as previously described(11, 16). Formation of the Fe-S cluster in IscU promoted by YtfE wild-type and mutants was evaluated in reaction mixtures containing apo-IscU (50 μM), IscS (4 μM), DTT (4 mM) and YtfE (150 μM), in 20 mM Tris-HCl pH 7.5 buffer supplemented with 150 mM NaCl. In all cases, addition of L-cysteine (3 mM) initiated the reaction performed under anaerobic conditions at room temperature for 1 h, and the process was analysed by visible spectroscopy in a Shimadzu UV-1700 spectrophotometer. Reaction mixtures that contained all components except YtfE served as controls.

### Crystallization and X-ray diffraction data collection and analysis

All the proteins used for crystallization procedures were purified in the buffer Tris-HCl 20 mM pH 7.5 supplemented with 150 mM NaCl and concentrated to 20 mg/mL. XRL hanging drop 24-well MD3-11 plates (Molecular Dimensions) were used for protein crystallization, with 0.5 mL of reservoir in each well. All experiments were done at room temperature.

Crystals of *E. coli* YtfE^M^ and YtfE^M^-E125L were prepared by the hanging drop vapour diffusion method using 1.0:0.8:0.2 μL mixtures of protein-reservoir-additive solutions. The reservoir solutions were constituted by Tris-HCl 0.1 M pH 7.5, 25 % PEG 4K and 0.2 M MgCl_2_ plus the additive NaCl (2 M) (Hampton Research). Crystals of YtfE^M^-E159L were prepared by the hanging drop vapour diffusion method using 1:1 μL mixtures of protein-reservoir solution, constituted by Tris-HCl 0.1 M pH 8.5, 30 % PEG 4K and 0.2 M MgCl_2_. These crystals were optimized using the micro-seeding technique (using crystals of YtfE^M^-E125L to obtain the first crystals) plus streak-seeding (using the first crystals of YtfE^M^-E159L).

Needle-shaped crystals were observed within a few minutes after plate set-up and grew to their maximal dimensions in 3-4 days. Crystals were harvested, immersed in a cryoprotectant solution with the same composition as the reservoir solution supplemented with 25% (v/v) glycerol, flash frozen in liquid nitrogen, and sent to a synchrotron beamline for data collection. The YtfE^M^ data set was collected at ALBA beamline XALOC(44) (Barcelona, Spain). The YtfE^M^-E125L data set was measured at Diamond Light Source, beamline I04 (DLS, Didcot, U.K.). The YtfE^M^-E159L data set was recorded at beamline ID30A-3 of the European Synchrotron Radiation Facility (ESRF, Grenoble, France). The data collection and processing statistics are shown in Supplemental Table 1.

The YtfE^M^ and YtfE^M^-E125L data sets were integrated and scaled with XDS(45), analysed with POINTLESS(46), and merged with AIMLESS(47). The YtfE^M^-E159L data set was integrated and scaled with XDS and AutoPROC(48), analysed with POINTLESS and scaled and merged with STARANISO(49) and AIMLESS. The structures were solved by molecular replacement with PHASER(50) via the CCP4(51) Graphics User Interface using the previously reported *E. coli* YtfE structure (PDB 5FNN)(12) as the search model. Following an initial refinement with REFMAC(52) the models were improved by sequential cycles of correction using COOT(53).

Refinement proceeded with PHENIX(54) in five macrocycle steps, with refinement of positional coordinates, individual isotropic atomic displacement parameters for all non-hydrogen atoms, occupancies and using non-crystallographic symmetry restraints for the two independent molecules in the asymmetric unit. Hydrogen atoms were added to the structural models and included in the refinement in calculated positions. The examination and editing of the models between refinements was done with COOT against σ_A_-weighted 2|F_o_| – |F_c_| and |F_o_| – |F_c_| electron density maps. TLS (translation-libration-screw) rigid body refinement of atomic displacement parameters was carried out for all structures, followed by refinement of individual isotropic B-factors. Two TLS groups were used for each chain, corresponding approximately to the two protein domains. Water molecules were added with PHENIX and verified with COOT. MOLPROBITY (55) was used to inspect the model geometry together with the validation tools available in COOT. The refinement statistics are included in Supplemental Table 2. Structure figures were created using the PyMOL(56) Molecular Graphics System, Version 2.3.4 Open Source (Schrödinger, LLC).

The tunnels were computed using the software MOLE 2.5(57), with default parameters (probe radius of 3 Å, interior threshold of 1.25 Å, minimum depth of 5 Å bottleneck radius of 1.25 Å) for the YtfE^M^ structure. For YtfE^M^-E159L the parameters were optimized (interior threshold of 1.18 Å and bottleneck radius of 1.13 Å). The indicated width of the tunnel at each point corresponds to the empty space between the Van der Waals spheres representing the atoms of the amino acid residues lining the tunnel.

The final atomic coordinates and experimental structure factors were deposited in the Worldwide Protein Data Bank(24) with the accession codes 7BHA, 7BHB and 7BHC for the YtfE^M^, YtfE^M^-E159L and YtfE^M^-E125L structures, respectively. Data collection and refinement statistics are summarised in Supplemental Tables 1 and 2.

## Acknowledgments

We would like to thank Lígia S. Nobre, Cláudia S. Freitas and Joana M. Baptista, former ITQB-NOVA members, for experimental support. We also thank Professor Miguel Teixeira of ITQB-NOVA for critical comments on the manuscript.

We thank the XALOC staff and floor coordinators at the synchrotron ALBA for the YtfE^M^ data collection. We acknowledge the ESRF for provision of synchrotron radiation facilities and we would like to thank Gianluca Santoni for assistance using the beamline ID30A-3 for the YtfE^M^-E159L data collection. Also, we would like to thank to Diamond Light Source for beamtime and the staff of beamline I04 for assistance with crystal testing and data collection of YtfE^M^-E125L.

This work was financially supported by Fundação para a Ciência e Tecnologia (Portugal) through fellowship SFRH/BD/118545/2016 (LSOS) and R&D unit LISBOA-01-0145-FEDER007660 (MostMicro) cofounded by FCT/MCTES and FEDER funds under the PT2020 Partnership Agreement. This work was partially supported by PPBI - Portuguese Platform of BioImaging (PPBI-POCI-01-0145-FEDER-022122) co-funded by national funds from OE - “Orçamento de Estado” and by European funds from FEDER - “Fundo Europeu de Desenvolvimento Regional”.

We also acknowledge funding from the European Union’s Horizon 2020 research and innovation programme under grant agreement No. 810856.

## Author Contributions

LSOS produced the proteins, performed biochemical experiments, protein-protein interactions studies and crystal production. CMR, PMM and LSOS collected and analysed the crystallographic data. LMS, LSOS and CMR wrote the manuscript with contributions from PMM. LMS supervised the work and designed the study.

## References

1. Overton TW, Justino MC, Li Y, Baptista JM, Melo AMP, Cole JA, Saraiva LM. 2008. Widespread distribution in pathogenic bacteria of di-iron proteins that repair oxidative and nitrosative damage to iron-sulfur centers. J Bacteriol 190:2004–13.

2. Justino MC, Almeida CC, Gonçalves VL, Teixeira M, Saraiva LM. 2006. *Escherichia coli* YtfE is a di-iron protein with an important function in assembly of iron-sulphur clusters. FEMS Microbiol Lett 257:278–84.

3. Justino MC, Baptista JM, Saraiva LM. 2009. Di-iron proteins of the Ric family are involved in iron-sulfur cluster repair. Biometals 22:99–108.

4. Justino MC, Vicente JB, Teixeira M, Saraiva LM. 2005. New genes implicated in the protection of anaerobically grown *Escherichia coli* against nitric oxide. J Biol Chem 280:2636–43.

5. Nobre LS, Meloni D, Teixeira M, Viscogliosi E, Saraiva LM. 2016. *Trichomonas vaginalis* Repair of Iron Centres Proteins: The Different Role of Two Paralogs. Protist 167:222–233.

6. Silva LO, Nobre LS, Mil-Homens D, Fialho A, Saraiva LM. 2018. Repair of Iron Centers RIC protein contributes to the virulence of *Staphylococcus aureus*. Virulence 9:312–317.

7. Harrington JC, Wong SMS, Rosadini C V., Garifulin O, Boyartchuk V, Akerley BJ. 2009. Resistance of *Haemophilus influenzae* to reactive nitrogen donors and gamma interferon-stimulated macrophages requires the formate-dependent nitrite reductase regulator-activated ytfE gene. Infect Immun 77:1945–1958.

8. Davis KM, Krupp J, Clark S, Isberg RR. 2019. Iron-Sulfur Cluster Repair Contributes to *Yersinia pseudotuberculosis* Survival within Deep Tissues. Infect Immun 87:e00533–19.

9. Nobre LS, Lousa D, Pacheco I, Soares CM, Teixeira M, Saraiva LM. 2015. Insights into the structure of the diiron site of RIC from *Escherichia coli*. FEBS Lett 589:426–431.

10. Todorovic S, Justino MC, Wellenreuther G, Hildebrandt P, Murgida DH, Meyer-Klaucke W, Saraiva LM. 2008. Iron-sulfur repair YtfE protein from *Escherichia coli:* Structural characterization of the di-iron center. J Biol Inorg Chem 13:765–770.

11. Nobre LS, Garcia-Serres R, Todorovic S, Hildebrandt P, Teixeira M, Latour J-M, Saraiva LM. 2014. *Escherichia coli* RIC is able to donate iron to iron-sulfur clusters. PLoS One 9:e95222.

12. Lo F-C, Hsieh C-C, Maestre-Reyna M, Chen C-Y, Ko T-P, Horng Y-C, Lai Y-C, Chiang Y-W, Chou C-M, Chiang C-H, Huang W-N, Lin Y-H, Bohle DS, Liaw W-F. 2016. Crystal Structure Analysis of the Repair of Iron Centers Protein YtfE and Its Interaction with NO. Chem - A Eur J 22:9768–9776.

13. Lu J, Bi B, Lai W, Chen H. 2019. Origin of Nitric Oxide Reduction Activity in Flavo–Diiron NO Reductase: Key Roles of the Second Coordination Sphere. Angew Chemie Int Ed 58:3795–3799.

14. Zheng L, Cash VL, Flint DH, Dean DR. 1998. Assembly of iron-sulfur clusters. Identification of an iscSUA-hscBA-fdx gene cluster from *Azotobacter vinelandii*. J Biol Chem 273:13264–13272.

15. Schwartz CJ, Djaman O, Imlay JA, Kiley PJ. 2000. The cysteine desulfurase, IscS, has a major role in in vivo Fe-S cluster formation in *Escherichia coli*. Proc Natl Acad Sci U S A 97:9009–9014.

16. Tokumoto U, Nomura S, Minami Y, Mihara H, Kato SS-i., Kurihara T, Esaki N, Kanazawa H, Matsubara H, Takahashi Y. 2002. Network of Protein-Protein Interactions among Iron-Sulfur Cluster Assembly Proteins in *Escherichia coli*. J Biochem 131:713–719.

17. Kim JH, Füzéry AK, Tonelli M, Ta DT, Westler WM, Vickery LE, Markley JL. 2009. Structure and Dynamics of the Iron-Sulfur Cluster Assembly Scaffold Protein IscU and Its Interaction with the Cochaperone HscB. Biochemistry 48:6062–6071.

18. Roche B, Aussel L, Ezraty B, Mandin P, Py B, Barras F. 2013. Iron/sulfur proteins biogenesis in prokaryotes: Formation, regulation and diversity. Biochim Biophys Acta - Bioenerg.

19. Blanc B, Gerez C, Ollagnier de Choudens S. 2015. Assembly of Fe/S proteins in bacterial systems. Biochemistry of the bacterial ISC system. Biochim Biophys Acta - Mol Cell Res 1853:1436–1447.

20. Karimova G, Pidoux J, Ullmann A, Ladant D. 1998. A bacterial two-hybrid system based on a reconstituted signal transduction pathway. Proc Natl Acad Sci U S A 95:5752–5756.

21. Kato SI, Mihara H, Kurihara T, Takahashi Y, Tokumoto U, Yoshimura T, Esaki N. 2002. Cys-328 of IscS and Cys-63 of IscU are the sites of disulfide bridge formation in a covalently bound IscS/IscU complex: Implications for the mechanism of iron-sulfur cluster assembly. Proc Natl Acad Sci U S A 99:5948–5952.

22. Kim JH, Bothe JR, Alderson TR, Markley JL. 2015. Tangled web of interactions among proteins involved in iron-sulfur cluster assembly as unraveled by NMR, SAXS, chemical crosslinking, and functional studies. Biochim Biophys Acta - Mol Cell Res.

23. Prischi F, Pastore C, Carroni M, Iannuzzi C, Adinolfi S, Temussi P, Pastore A. 2010. Of the vulnerability of orphan complex proteins: The case study of the *E. coli* IscU and IscS proteins. Protein Expr Purif 73:161–166.

24. Burley SK, Berman HM, Bhikadiya C, Bi C, Chen L, Costanzo L Di, Christie C, Duarte JM, Dutta S, Feng Z, Ghosh S, Goodsell DS, Green RK, Guranovic V, Guzenko D, Hudson BP, Liang Y, Lowe R, Peisach E, Periskova I, Randle C, Rose A, Sekharan M, Shao C, Tao YP, Valasatava Y, Voigt M, Westbrook J, Young J, Zardecki C, Zhuravleva M, Kurisu G, Nakamura H, Kengaku Y, Cho H, Sato J, Kim JY, Ikegawa Y, Nakagawa A, Yamashita R, Kudou T, Bekker GJ, Suzuki H, Iwata T, Yokochi M, Kobayashi N, Fujiwara T, Velankar S, Kleywegt GJ, Anyango S, Armstrong DR, Berrisford JM, Conroy MJ, Dana JM, Deshpande M, Gane P, Gáborová R, Gupta D, Gutmanas A, Koča J, Mak L, Mir S, Mukhopadhyay A, Nadzirin N, Nair S, Patwardhan A, Paysan-Lafosse T, Pravda L, Salih O, Sehnal D, Varadi M, Vǎreková R, Markley JL, Hoch JC, Romero PR, Baskaran K, Maziuk D, Ulrich EL, Wedell JR, Yao H, Livny M, Ioannidis YE. 2019. Protein Data Bank: The single global archive for 3D macromolecular structure data. Nucleic Acids Res 47:D520–D528.

25. Brunskill EW, De Jonge BLMM, Bayles KW. 1997. The *Staphylococcus aureus* scdA gene: A novel locus that affects cell division and morphogenesis. Microbiology 143:2877–2882.

26. Chang W, Small DA, Toghrol F, Bentley WE. 2006. Global transcriptorne analysis of *Staphylococcus aureus* response to hydrogen peroxide. J Bacteriol 188:1648–1659.

27. Alvarez-Carreño C, Alva V, Becerra A, Lazcano A. 2018. Structure, function and evolution of the hemerythrin-like domain superfamily. Protein Sci 27:848–860.

28. Layer G, Ollagnier-De Choudens S, Sanakis Y, Fontecave M. 2006. Iron-sulfur cluster biosynthesis: Characterization of *Escherichia coli* CyaY as an iron donor for the assembly of [2Fe-2S] clusters in the scaffold IscU. J Biol Chem 281:16256–16263.

29. Skovran E, Lauhon CT, Downs DM. 2004. Lack of YggX results in chronic oxidative stress and uncovers subtle defects in Fe-S cluster metabolism in *Salmonella enterica*. J Bacteriol 186:7626–7634.

30. Ding H, Clark RJ. 2004. Characterization of iron binding in IscA, an ancient iron-sulphur cluster assembly protein. Biochem J 379:433–440.

31. Lu J, Yang J, Tan G, Ding H. 2008. Complementary roles of SufA and IscA in the biogenesis of iron-sulfur clusters in *Escherichia coli*. Biochem J 409:535–543.

32. Silva LSO, Baptista JM, Batley C, Andrews SC, Saraiva LM. 2018. The Di-iron RIC Protein (YtfE) of *Escherichia coli* Interacts with the DNA-Binding Protein from Starved Cells (Dps) To Diminish RIC Protein-Mediated Redox Stress. J Bacteriol 200:e00527–18.

33. Tatur J, Hagen WR, Matias PM. 2007. Crystal structure of the ferritin from the hyperthermophilic archaeal anaerobe *Pyrococcus furiosus*. J Biol Inorg Chem 12:615–630.

34. Macedo S, Romão C V., Mitchell E, Matias PM, Liu MY, Xavier A V., LeGall J, Teixeira M, Lindley P, Carrondo MA. 2003. The nature of the di-iron site in the bacterio-ferritin from *Desulfovibrio desulfuricans*. Nat Struct Biol 10:285–290.

35. Swartz L, Kuchinskas M, Li H, Poulos TL, Lanzilotta WN. 2006. Redox-dependent structural changes in the *Azotobacter vinelandii* bacterioferritin: New insights into the ferroxidase and iron transport mechanism. Biochemistry 45:4421–4428.

36. Weeratunga SK, Lovell S, Yao H, Battaile KP, Fischer CJ, Gee CE, Rivera M. 2010. Structural studies of bacterioferritin B from *Pseudomonas aeruginosa* suggest a gating mechanism for iron uptake via the ferroxidase center. Biochemistry 49:1160–1175.

37. Rui H, Rivera M, Im W. 2012. Protein dynamics and ion traffic in bacterioferritin. Biochemistry 51:9900–9910.

38. Ebrahimi KH, Hagedoorn P-L, Hagen WR. 2015. Unity in the biochemistry of the iron-storage proteins ferritin and bacterioferritin. Chem Rev 115:295–326.

39. Roche B, Huguenot A, Barras F, Py B. 2015. The iron-binding CyaY and IscX proteins assist the ISC-catalyzed Fe-S biogenesis in *Escherichia coli*. Mol Microbiol 95:605–23.

40. Vinella D, Brochier-Armanet C, Loiseau L, Talla E, Barras F. 2009. Iron-Sulfur (Fe/S) Protein Biogenesis: Phylogenomic and Genetic Studies of A-Type Carriers. PLoS Genet 5:e1000497.

41. Tan G, Lu J, Bitoun JP, Huang H, Ding H. 2009. IscA/SufA paralogues are required for the [4Fe-4S] cluster assembly in enzymes of multiple physiological pathways in *Escherichia coli* under aerobic growth conditions. Biochem J 420:463–472.

42. Wilson CGM, Magliery TJ, Regan L. 2004. Detecting protein-protein interactions with GFP-fragment reassembly. Nat Methods 1:255–262.

43. Fischer DS, Price DC. 1964. A Simple Serum Iron Method Using the New Sensitive Chromogen Tripyridyl-s-triazine. Clin Chem 10:21–31.

44. Juanhuix J, Gil-Ortiz F, Cuní G, Colldelram C, Nicolás J, Lidón J, Boter E, Ruget C, Ferrer S, Benach J. 2014. Developments in optics and performance at BL13-XALOC, the macromolecular crystallography beamline at the Alba Synchrotron. J Synchrotron Radiat 21:679–689.

45. Kabsch W. 2010. XDS. Acta Crystallogr D Biol Crystallogr 66:125–32.

46. Evans P. 2006. Scaling and assessment of data quality, p. 72–82. In Acta Crystallographica Section D: Biological Crystallography. International Union of Crystallography.

47. Evans PR, Murshudov GN. 2013. How good are my data and what is the resolution? Acta Crystallogr Sect D Biol Crystallogr 69:1204–1214.

48. Vonrhein C, Flensburg C, Keller P, Sharff A, Smart O, Paciorek W, Womack T, Bricogne G. 2011. Data processing and analysis with the autoPROC toolbox. Acta Crystallogr Sect D Biol Crystallogr 67:293–302.

49. Tickle, I.J., Flensburg, C., Keller, P., Paciorek, W., Sharff, A., Vonrhein, C., Bricogne G. 2018. STARANISO. Cambridge, United Kingdom Glob Phasing Ltd.

50. McCoy AJ. 2006. Solving structures of protein complexes by molecular replacement with Phaser, p. 32–41. In Acta Crystallographica Section D: Biological Crystallography. International Union of Crystallography.

51. Potterton E, Briggs P, Turkenburg M, Dodson E. 2003. A graphical user interface to the CCP4 program suite. Acta Crystallogr - Sect D Biol Crystallogr 59:1131–1137.

52. Murshudov GN, Skubák P, Lebedev AA, Pannu NS, Steiner RA, Nicholls RA, Winn MD, Long F, Vagin AA. 2011. REFMAC5 for the refinement of macromolecular crystal structures. Acta Crystallogr Sect D Biol Crystallogr 67:355–367.

53. Emsley P, Lohkamp B, Scott WG, Cowtan K. 2010. Features and development of Coot. Acta Crystallogr Sect D Biol Crystallogr 66:486–501.

54. Adams PD, Afonine P V., Bunkóczi G, Chen VB, Davis IW, Echols N, Headd JJ, Hung LW, Kapral GJ, Grosse-Kunstleve RW, McCoy AJ, Moriarty NW, Oeffner R, Read RJ, Richardson DC, Richardson JS, Terwilliger TC, Zwart PH. 2010. PHENIX: A comprehensive Python-based system for macromolecular structure solution. Acta Crystallogr Sect D Biol Crystallogr 66:213–221.

55. Chen VB, Arendall WB, Headd JJ, Keedy DA, Immormino RM, Kapral GJ, Murray LW, Richardson JS, Richardson DC. 2010. MolProbity: All-atom structure validation for macromolecular crystallography. Acta Crystallogr Sect D Biol Crystallogr 66:12–21.

56. DeLano WL. 2020. The PyMOL Molecular Graphics System, Version 2.3. Schrödinger LLC.

57. Sehnal D, Vařeková RS, Berka K, Pravda L, Navrátilová V, Banáš P, Ionescu CM, Otyepka M, Koča J. 2013. MOLE 2.0: Advanced approach for analysis of biomacromolecular channels. J Cheminform 5:39.

58. Baker NA, Sept D, Joseph S, Holst MJ, McCammon JA. 2001. Electrostatics of nanosystems: Application to microtubules and the ribosome. Proc Natl Acad Sci U S A 98:10037–10041.

59. Dolinsky TJ, Nielsen JE, McCammon JA, Baker NA. 2004. PDB2PQR: An automated pipeline for the setup of Poisson-Boltzmann electrostatics calculations. Nucleic Acids Res 32:W665–W667.

